# Characterization of mitotic phenotypes associated with a MYC synthetic lethal compound

**DOI:** 10.1101/2023.04.03.535438

**Authors:** Qiong Shi, Xuemei Liu, Julia Kalashova, Chenglu Yang, Hongmei Li, Yan Long, Ting Zhang, Shenqiu Zhang, Gang Lv, Jian Huang, Hong Liu, Jing Zhang, Thaddeus D. Allen, Dun Yang

## Abstract

Therapeutic targeting of MYC directly has proven difficult, but several means to target MYC indirectly using a synthetic lethal drug approach have been proposed. Synthetic lethal approaches for MYC have sought to take advantage of vulnerabilities MYC imposes related to either metabolic reprogramming, apoptotic signaling or the cycling of cancer cells. Here, we describe in detail the cell division phenotypes induced by a MYC synthetic lethal compound, dimethylfasudil (diMF). DiMF is a known ROCK inhibitor, but structurally related ROCK inhibitors are not synthetic lethal with MYC, so the activity of diMF is not related to blockade of this family of kinases. Instead, this compound induced multiple cell cycle-related liabilities. These included the early mitotic arrest of cycling cells followed by mitotic catastrophe-induced death and the induction of polyploidy in cells that do manage to pass through mitosis. As early as prometaphase, we noted diminished staining for BUB1 kinase, which binds to kinetochores and regulates the mitotic spindle checkpoint and chromosome congression. Kinetochore proteins, such as CENP-F, failed to localize at the metaphase plate, confirming a deficit in centromere assembly. This, presumably, contributed to the development of segregation anomalies in diMF-treated cells. In anaphase cells, the protein regulator of cytokinesis 1 (PRC1), failed to be recruited to the midzone, leading to a cascade of defects that included failed recruitment of the chromosomal passenger protein complex, the centralspindlin complex and polo-like-kinase 1 (PLK1). These observations correlate well with the cell death phenotypes induced by diMF, which may serve as a prototype MYC synthetic lethal compound to explore synthetic lethal therapy or as a scaffold upon which to build superior compounds. The phenotypes described here serve as examples of MYC synthetic lethal drug effects that can be used to explore and maximize drug discovery programs.

## Introduction

*MYC* is the prototype member of a family of mammalian genes. It was first cloned as a homolog of *v-Myc*, the avian viral myelocytomatosis oncogene (Vennstrom et al., 1982). Two additional genes, *MYCN* (for neuroblastoma-derived homolog) and *MYCL* (lung carcinoma-derived homolog), complete this proto-oncogene family and together these genes encode what may be the most widely deregulated oncoproteins in human cancer (Dhanasekaran et al., 2022). They encode transcription factors with a C-terminal basic region, a helix-loop-helix and a leucine zipper (bHLHZip) domain that dimerizes at DNA binding sites in concert with a second transcription factor, MAX. However, MYC also interacts with the basal transcriptional machinery and recruits higher order chromatin regulatory complexes to influence transcription, and this does not necessarily require direct DNA binding (Lourenco et al., 2021). In addition, a cleavage product of full-length MYC, known as MYC-nick, possesses non-nuclear functions that can contribute to carcinogenesis (Conacci-Sorrell et al., 2014).

A therapy that inhibits MYC has long been a goal for both academic and industry research (Dhanasekaran et al., 2022; Lourenco et al., 2021), but thus far, direct inhibition of MYC has proven difficult. An alternative means to target MYC expressing cells is to develop therapies that are synthetic lethal with MYC overexpression. Synthetic lethality can occur when the same oncogenic drivers that promote carcinogenesis, also introduce vulnerabilities to cancer cells. Targeting these vulnerabilities can result in enhanced anticancer activity in vulnerable cells, while leaving normal cells, lacking oncogenic drivers, healthy. Known synthetic lethal targets for MYC overexpressing cells include AURKB (Diaz et al., 2015; Yang et al., 2010), CDK1 (Goga et al., 2007), the serine-threonine kinase PIM1 (Horiuchi et al., 2016) and TNFRSF10B (Wang et al., 2004).

Dimethylfasudil (diMF) is a chemical compound that has been shown to enhance killing of cells that overexpress MYC (Zhang et al., 2021). Here, we have expanded these observations to include assays of 74 human cancer cell lines and the analysis of xenografts expressing varying amounts of MYC. We can now definitively conclude that diMF is a MYC synthetic lethal compound. Although diMF is an analog of clinically used ROCK inhibitors, the synthetic lethal killing of MYC expressing cells is not related to ROCK inhibition. We hypothesized that the mode-of-action of diMF to effect synthetic lethality might be through inhibition of AURKB, since the phenotype induced by diMF mimics that induced by AURKB inhibitors in our experimental cell lines. However, this proved not to be the case. Instead, diMF appears to limit the mitotic localization of key proteins that regulate chromosome segregation and cytokinesis. This correlates with a diminished mitotic spindle, kinetochore dysfunction and an anaphase midzone organization defect that prevents midbody formation and successful cytokinesis. DiMF represents a new class of synthetic lethal compound that disrupts mitotic progression through the mis-localization of proteins that direct mitosis, rather than through the direct inhibition of AURK catalytic activity or through the blockade of spindle microtubule dynamics.

## Materials and Methods

### Chemicals and Reagents

The commercial source of dimethylfausadil (diMF), also known as H-1152 dihydrochloride (CAS 451462-58-1), was Santa Cruz Biotechnology (Dallas, TX). Fasudil (Cat. # 0541/10) and hydroxyfasudil hydrochloride (Cat. #2415) were purchased from Tocris Bioscience (Bristol, UK). Compound 7 (IUPAC name: 5-(1,4-Diazepan-1-ylsulfonyl)-4-methyl-isoquinoline) was synthesized in-house and structure and purity were confirmed analytically. Ripasudil K-115 (Cat. #S7995) and the AURK inhibitors barasertib (AZD1152-HQPA, Cat. #S1147) and alisertib (MLN8237, Cat. #S1133) were purchased from Selleck Chemicals (Houston, TX). Slide mounting media containing 4′,6-Diamidino-2-phenylindole dihydrochloride (DAPI) was purchased from Vector Laboratories (Burlingame, CA, Cat. #H-1150) and the Yeasen Institute of Biotechnology (Shanghai, China, Cat. #36308ES11).

### Cell Lines and Culture Conditions

Human cancer cell lines were purchased from the American Type Culture Collection (ATCC, Manassas, VA). Derivation of experimental retinal pigment epithelial (RPE) cell lines RPE-MYC and RPE-MYC^BCL2^ have been previously described (Goga et al., 2007; Yang et al., 2010; Zhang et al., 2020a). Briefly, RPE-MYC cells were generated by transfecting hTERT-immortalized human primary retinal pigment epithelial cells (RPE-hTERT) with a vector expressing the neomycin resistance gene alongside the MYC oncogene, followed by selection with 800 μg/ml G418 (Geneticin, Thermo-Fisher Scientific, Waltham, MA). RPE-MYC cells were transfected with a construct harboring BCL2 and selected with 1 μg/ml of puromycin (Thermo-Fisher) to generate RPE-MYC^BCL2^ cells. Isogenic RPE cell lines were cultured in DMEM supplemented with 10% fetal bovine serum and antibiotics, at 5% CO_2_ in a humidified incubator. Human cancer cells were cultured in media as shown in Supplementary Table 1. To ensure that cell lines were free of mycoplasma contamination, a Universal Mycoplasma Detection Kit (ATCC1 30-1012K™) was used to assay select cell lines for the mycoplasma 16S rRNA-coding region after experimentation.

### Viability Assays

Trypan blue exclusion assays were used to determine cell viability. Cells were plated into 6-well plates and allowed to adhere overnight at 37°C before being incubated with a vehicle control or drug for indicated times. Cells were washed with PBS once, treated with a 0.25% trypsin-EDTA solution to free cells from the plate and then resuspended in DMEM. After staining with trypan blue (0.4%, 1:1) (Sangon, Cat. #E607338, Shanghai, China), cells were counted with a hematocytometer under a high powered microscope or using a Luna Cell Counter (Logos Biosystems, Inc., South Korea). Viability was expressed as the percent dead cells in the population. Data is presented from three independent experiments that were performed with sequentially passaged cells, and each experiment was done in triplicate (n = 9).

### Western Analysis

Whole cell protein extracts were prepared by treating adherent cells for 15 minutes at 4°C with lysis buffer (50 mM Tris at pH 7.5, 200 mM NaCl, 0.1% SDS, 1% Triton X-100, 0.1 mM DTT, and 0.5 mM EGTA) that was supplemented with BD BaculoGold Protease Inhibitor Cocktail (BD Biosciences, San Jose, CA, Cat. # 554779). Extracts were cleared by centrifugation at 8,000×g for 10 minutes. Final protein concentration was determined using the Bio-Rad Protein Assay reagent (Hercules, CA, Cat. # 5000006). For analysis, 50 μg of protein were resolved using a NuPAGE (4– 12%) Bis-Tris gel (Thermo-Fisher Scientific, Cat. # NP0322PK2). After the run, protein was transferred to a nitrocellulose membrane (Bio-Rad, Cat. # 1620150). Membranes were blocked with 5% non-fat milk in PBS for 1 hour at room temperature before a 4°C overnight incubation with primary antibody diluted in blocking buffer. Primary antibodies used for Westerns were antibodies for MYC (Cat. # ab32072) or MYCN (Cat. # ab24293), both from Abcam (Cambridge, UK), a β-Tubulin antibody from Millipore Sigma (Burlington, MA, Cat. # T4026) and an antibody recognizing MYCL from Cell Signaling Technology (Danvers, MA, Clone Cat. # 76266) Detection was with horseradish peroxidase (HRP)-conjugated mouse anti-rabbit IgG (1:5000, Santa Cruz Biotechnology, Cat. # sc-2357). Visualization was with a SuperSignal West Femto ECL detection kit (Thermo-Fisher Scientific, Cat. # 34095).

### Immunofluorescent Staining

Immunofluorescent staining experiments were performed as previously described (Yang et al., 2010). RPE-MYC^BCL2^ cells were cultured on coverslips coated with 0.1% gelatin in a six-well plate and allowed to adhere overnight at 37°C. After drug treatment, cells were washed with PBS, fixed with 4% paraformaldehyde (PFA) for 10 minutes in the presence of 0.5% TritonX-100, and then incubated with 5% BSA as a blocking buffer for 1 hour at room temperature. Immunofluorescent staining procedures were typically performed with incubation of the primary antibody for 2 hours at room temperature and then with a fluorescent probe-conjugated secondary antibody for 1 hour at room temperature. Cells were mounted on microscope slides with DAPI-containing Vectashield and observed with a 63X/1.4 N.A. oil objective or an EVOS FL Auto microscope (Thermo-Fisher Scientific).

### Antibodies for Immunofluorescent Staining

To detect AURKB, we utilized a rabbit monoclonal antibody from Cell Signaling (Cat. #3094) or a mouse monoclonal AURKB antibody (Cat. # A78720) from Transduction Laboratories (now BD Bioscience, San Jose, CA). Rabbit monoclonal antibody (Cat. #1173-1) and mouse monoclonal antibody (Cat. #05-806) recognizing Histone 3 phosphorylated at Ser10, were obtained from Epitomics (now Abcam) and Upstate Biotechnology (Lake Placid, NY), respectively. Murine monoclonal antibodies recognizing β-tubulin (Cat. #T4026) and γ-tubulin (Cat. #T6557), were from Millipore Sigma (Burlington, MA). A rabbit polyclonal inner centromere protein (INCENP) antibody was gifted (T. Stukenberg, University of Virginia, Charlottesville, and W. C. Earnshaw, University of Edinburgh), as was CREST serum (B. R. Brinkley, Baylor College of Medicine, Houston). Goat PRC1 antibody (K-18, Cat. #sc-9342, discontinued) and rabbit polyclonal to MKLP1 (N-19, Cat. #sc-867, discontinued) were purchased from Santa Cruz Biotechnology. Mouse monoclonal PLK1 antibody (PL6/PL2 Cat. #33-1700) was from Thermo-Fisher Scientific. A rabbit polyclonal antibody recognizing the protein Ninein/NIN was obtained from Abcam (Cat. #ab4447). Bub1 mouse monoclonal antibody (Cat. #MAB3610) was obtained from Chemicon (now Millipore Sigma). Rabbit polyclonal antibody recognizing the Kif11/Eg5 protein (Cat. #AKIN03, discontinued) was purchased from Cytoskeleton, Inc. (Denver, CO). Rabbit polyclonal antibody recognizing CENP-F (Cat. #NB500-101) was purchased from Novus Biologics (Littleton, CO). Secondary antibodies used included Rhodamine Red™-X (RRX) AffiniPure Donkey Anti-Human IgG (H+L) (Cat. # 709-295-149), Rhodamine Red™-X (RRX) AffiniPure Bovine Anti-Goat IgG (H+L) (Cat. # 805-295-180), Fluorescein (FITC) AffiniPure Goat Anti-Rabbit IgG (H+L) (Cat. # 111-095-003), Rhodamine (TRITC) AffiniPure Goat Anti-Rabbit IgG (H+L) (Cat. # 111-025-003) and Alexa Fluor^®^ 488 AffiniPure Goat Anti-Mouse IgG (H+L) (Cat. # 115-545-003), all obtained from Jackson ImmunoResearch Laboratories (West Grove, PA). A Rhodamine (TRITC)–conjugated Goat Anti-Mouse IgG(H+L) was purchased from Proteintech Group, Inc. (Wuhan, China).

### Murine Xenograft Experiments

The human small cell lung cancer cell lines NCI-H524, NCI-H841 and NCI-H211 and the non-small cell lung carcinoma NCI-H2199 were grown in the flanks of NCG triple-immunodeficient mice (NOD-Prkd^cem26Cd52^Il2rg^em26Cd22^/NjuCrl, strain T0014575, Nanjing Ji Cui Yao Kang Biology Science And Technology Co.,Ltd., China). One million to ten million cells were used per injection. When tumors reached approximately 50-70 mm^3^, tumor-bearing mice were randomized to receive oral gavage with either vehicle or diMF (60 mg/kg body weight for each injection), twice daily. For each group, n=10 mice. Tumors were measured using calipers every 3 days until the end of the experiment. Tumor volume was calculated using the formula, Vol. = (Width(2) × Length)/2.

### Statistical Analysis

Statistical analysis was carried out using Prism 9 software (GraphPad Software, San Diego, CA). For viability assays, experiments were carried out in triplicate and data points are presented as an average with error bars as the standard deviation.

## Results

### Dimethylfasudil (diMF) is a MYC Synthetic Lethal Compound

The compound diMF was previously identified in a screen of clinically used compounds and their close analogs as able to induce transient cell cycle arrest, mitotic catastrophe and polyploidy (Zhang et al., 2021). In this high-content screen, compounds were selected for their ability to induce these phenotypes in retinal pigment epithelial (RPE) cells that ectopically expressed the MYC oncogene (RPE-MYC), while leaving a non-transformed counterpart (RPE-neo) viable and healthy. It was confirmed that MYC-induced sensitization to diMF was not a general effect of oncogenic transformation. Expression of other oncoproteins, like activated RAS, RAF or dominant negative p53, did not sensitize to diMF. Likewise, MYC did not have the generalized effect of sensitizing RPE cells to all apoptotic stimuli, as apoptosis induced by a panel of chemotherapeutic drugs was not influenced by MYC (Zhang et al., 2021).

Several studies now describe synthetic lethality between an experimental drug treatment and MYC expression (Goga et al., 2007; Horiuchi et al., 2016; Yang et al., 2010). While screening efforts are often carried out in isogenic cell lines, confirmation studies are usually carried out in a select few human cell lines and murine models (Cermelli et al., 2014). Before designating diMF as a MYC synthetic lethal compound we wished to carry out viability studies in a much larger panel of human cancer cell lines expressing a range of MYC protein. We assayed MYC protein levels in 74 human cancer cell lines individually by Western analysis (Table S1). The intensity of the MYC protein band was normalized to β-tubulin as a housekeeping, loading control protein to produce a metric of MYC protein level in each line. The MYC^High^ group also includes eight cell lines with abundant MYCN and two cell lines abundantly expressing MYCL. This analysis resulted in the division of cell lines into high (n=51, 69% of cell lines) and low (n=23, 31%) expression groups. Cell death with diMF (6 μM) was assayed using a trypan blue exclusion assay following treatment of each cell line for 24 hours. Ranking the amount of cell death induced by diMF revealed a pattern between the high and low MYC expression groups (Figure 1A) with 27 of the top 28 and 42 of the top 50 cell lines being found in the MYC^High^ group. The difference in response of the MYC^High^ and MYC^Low^ groups was statistically significant (Figure 1B, *p<0.0001*).

**Figure 1:**
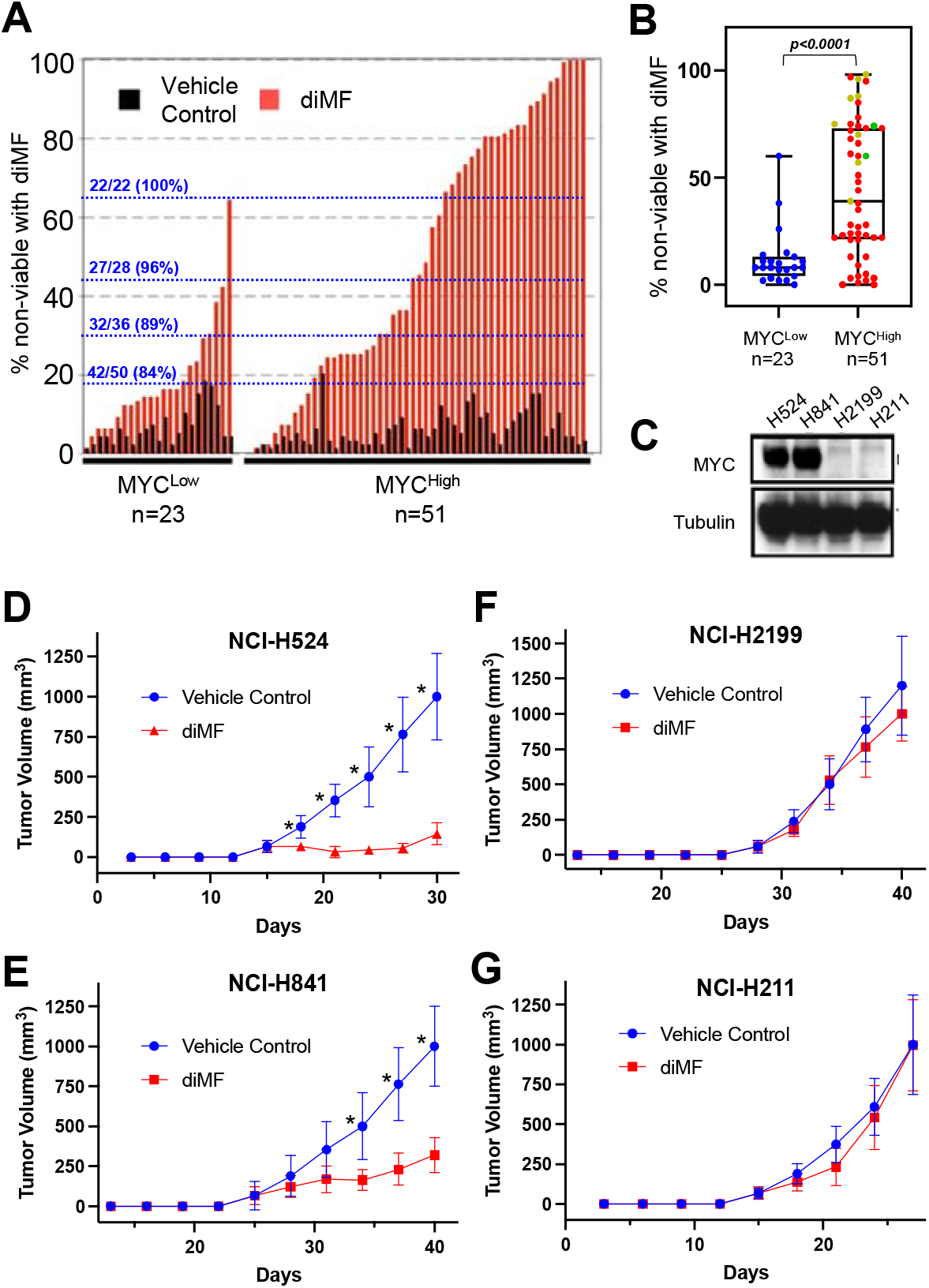
MYC expression predicts the sensitivity to diMF across a wide range of cancer cell lines. **(A)** The trypan blue exclusion assay was used to quantify the percentage of dead cells. The sensitivity of 74 human cancer cell lines to diMF (6 μM, red bars) was ranked following 4 days (96 hours) of exposure. The percent cell death with vehicle (0.1% DMSO) is included as black bars. Data is the average of 2 experiments with 3 replicates (n=6). Cells were grouped according to MYC abundance with 51 cell lines (69%) having abundant MYC and 23 (31%) having low MYC. The ratio of cell lines falling in the group with abundant MYC with a percent cell death above the dotting line is shown in blue. **(B)** Range in the percent non-viable cells for the 74 cell lines. Red, yellow, and green dots represent cell lines with abundant MYC, MYCN and MYCL, respectively. The *p-value* is from a Mann-Whitney U Test. **(C)** Western analysis of MYC in the small cell lung cancer cell lines NCI-H524, NCI-H841 and NCI-H211 and the non-small cell lung carcinoma NCI-H2199. Tubulin was used as a loading control. **(D-G)** Growth curves for xenografts of **(D)** NCI-H524, **(E)** NCI-H841, **(F)** NCI-H2199 and **(G)** NCI-H211, grown in immunocompromised mice. Treatment was twice daily by oral gavage with vehicle control (0.1% DMSO) or diMF (60 mg/kg body weight). For each group in **(D-G)**, n=10 mice. * is a *p-value < 0.001*, resulting from an unpaired t-test with Welch correction.

To determine if these *in vitro* observations on the ability to induce cell death translate into differences in the *in vivo* anticancer activity of diMF, we selected two MYC expressing (NCI-H524, NCI-H841) and two low MYC (NCI-H2199, NCI-H211) cell lines (Figure 1C) to assay diMF in xenograft models. The response of diMF *in vivo* was pronounced in the MYC expressing xenografts (Figure 1D, E), while the response was blunted in the xenografts where MYC was low (Figure 1F, G). We concluded from these confirmatory studies that diMF is indeed a MYC synthetic lethal compound. Since our cell line panel contains cancer cells from multiple indications, we also concluded that MYC expression, not tissue-of-origin is the most important determinant of sensitivity.

### MYC synthetic lethality with dimethylfasudil (diMF) is not dependent on ROCK inhibition

To search for elements in the diMF molecule that are determinants of its activity, we took advantage of the fact that many similar molecules have been described as inhibitors of ROCK kinases. IC_50_ values for known ROCK inhibitors and one related molecule, compound 7, are listed in Table 1 for the inhibition of ROCK1 and ROCK2 kinase activity. All share a similar structure with an isoquinoline ring attached via a sulfonamide linkage with the aromatic nitrogen of a hyperpiperazine ring. The compounds differ from diMF in the location of two distinct methyl groups, one on the hyperpiperazine ring and one on the isoquinoline ring.

**Table 1:**
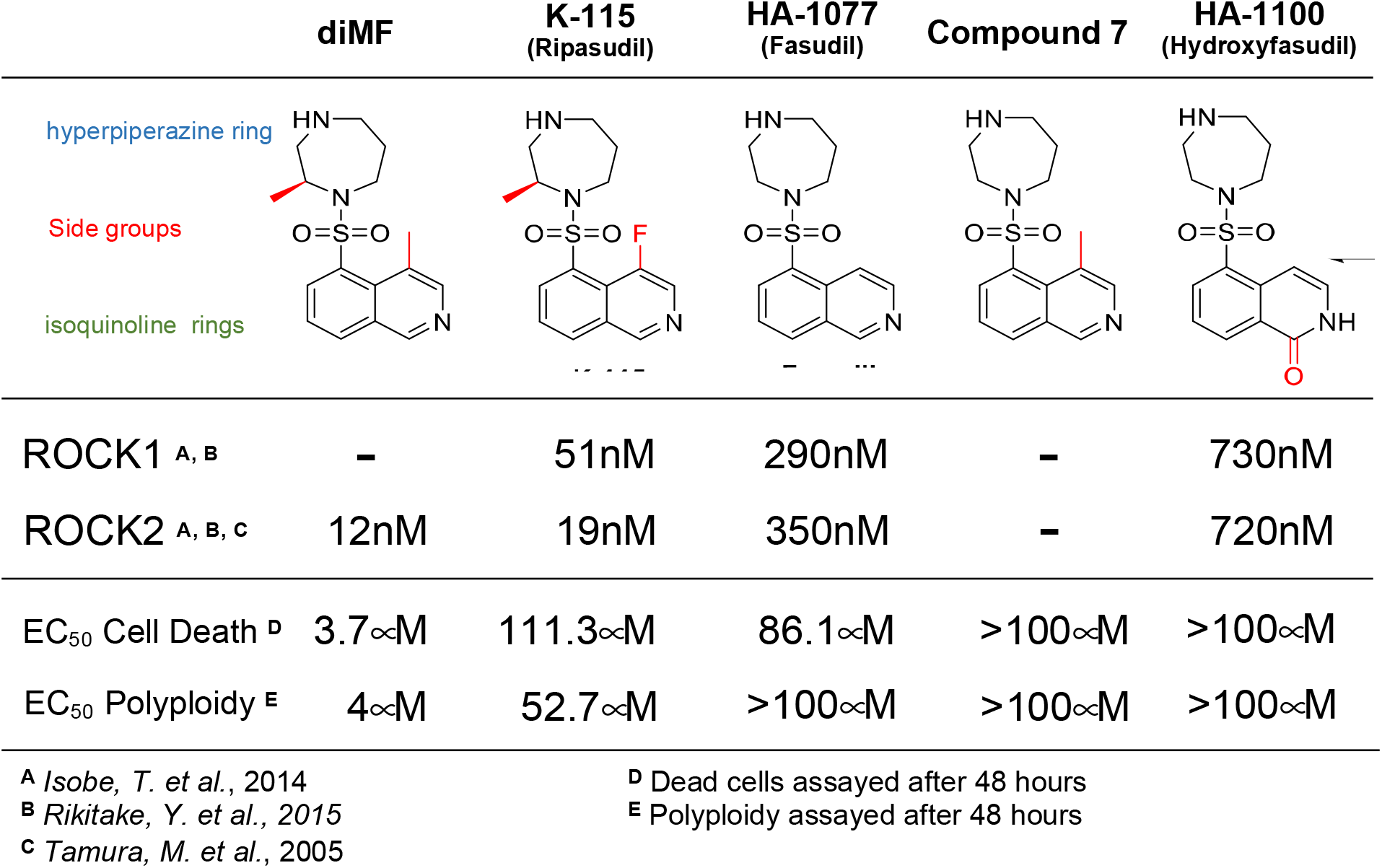
Comparison of EC_50_ for ROCK1,2 inhibition with that found for defective cell division phenotypes in RPE-MYC^BCL2^ cells

The ability to induce cell death (Figure S1A) and the ability to induce polyploid cells (Figure S1B) were both compromised by the absence of either one of the two methyl groups. The EC_50_ for both the induction of apoptosis after 48 hours of diMF exposure and the formation of polyploid cells after 48 hours, were substantially higher for these compounds than for diMF (Table 1). There was, therefore, no relationship between the cell division phenotypes induced in RPE-MYC^BCL2^ cells and ROCK inhibitory activity. Since the analogs used here differ only in the presence or absence of one or both of the two distinct methyl groups (highlighted red in Table 1), we also conclude that these are functional groups absolutely required for diMF to induce cell division phenotypes.

### Dimethylfasudil (diMF) does not inhibit catalytic activity of AURKs

Previously, surrogate markers for AURK activity were used to examine if diMF could target this kinase family, but this was carried out only using the experimental cell line, RPE-MYC (Zhang et al., 2021). Mitotic phosphorylation of Histone 3 at serine 10 (H3PS10), a surrogate for AURKB catalytic activity (Hsu et al., 2000; Li et al., 2021; Yan et al., 2022) and the autophosphorylation of AURKA at Thr288, a surrogate for its activity (Dauch et al., 2016; Hirota et al., 2003), were unaltered by diMF treatment in RPE-MYC cells. Here we examined the surrogate marker H3PS10 further with the goal of determining if the findings were a product of our experimental RPE cells, or more generalizable. In a variety of human cancer cell lines, H3PS10 was abundant in cells arrested in prometaphase after diMF treatment (Figure S2), confirming that with the surrogate marker assay, the catalytic activity of AURKB does not appear to be the target of diMF. Despite this, diMF induced polyploidy in all cell human cancer cell lines examined, confirming activity of the compound.

As an additional functional assay of AURK inhibition early in mitosis, we examined centrosome number in diMF treated cells. Inhibitors of AURKs are able to induce amplification of centrosomes and this occurs in cells arrested at prometaphase. Nucleation of the amplified centrosomes occurs, resulting in multipolar spindle formation (Faisal et al., 2011; Yang et al., 2010). We have previously found that even partial inhibition of AURK activity with naturally occurring phytochemicals can induce centrosome amplification in RPE-MYC^BCL2^ cells (Li et al., 2021; Yan et al., 2022), making detection of this phenotype a very sensitive assay for diminished AURK catalytic activity. Immunofluorescent staining for the centrosome marker γ-tubulin (Figure S3A) did not reveal increased centrosome number. We confirmed this with staining for the protein Ninein, which is required for maturation of centrosomes (Ou et al., 2002). We did not observe an abnormal number of Ninein foci (Figure S3B). We concluded from these observations that centrosome amplification is not significantly enhanced by diMF, consistent with a mechanism of action that is distinct from inhibition of AURK catalytic activity.

### Diminished mitotic spindle microtubule density

The use of spindle toxins can predispose to the formation of polyploidy (Saini et al., 2022; Zhang et al., 2014). Given this overlapping phenotype with diMF, we next examined the integrity of the mitotic spindle in diMF-treated cells. We immunohistochemically stained cells with β-tubulin, visualizing the spindle. Staining was suggestive of a disorganized and diminished spindle structure (Figure 2A). This was most pronounced as cells progressed to anaphase. The poles in anaphase cells, the centrosome-mediated microtubular organizing centers, were still discernable, but not the polar microtubules of the spindle itself. This suggested a lack of microtubule density. Immunofluorescent staining for the motor protein Kif11 (also known as Eg5) confirmed this finding (compare metaphase cells, Figure 2B). Kif11 is a plus-end-directed kinesin that crosslinks and slides antiparallel microtubules apart to maintain spindle bipolarity (Kapitein et al., 2005). This function is enabled though homotetramerization which places two ATPase head groups at each end, facilitating microtubule bundling as well as sustained outward forces to push the spindle poles apart. In RPE-MYC^BCL2^ cells, staining for Kif11 was disorganized and diminished with diMF treatment. Overall, this immunostaining data suggests a weakened spindle that lacks the microtubule polymer density found in control cells.

**Figure 2:**
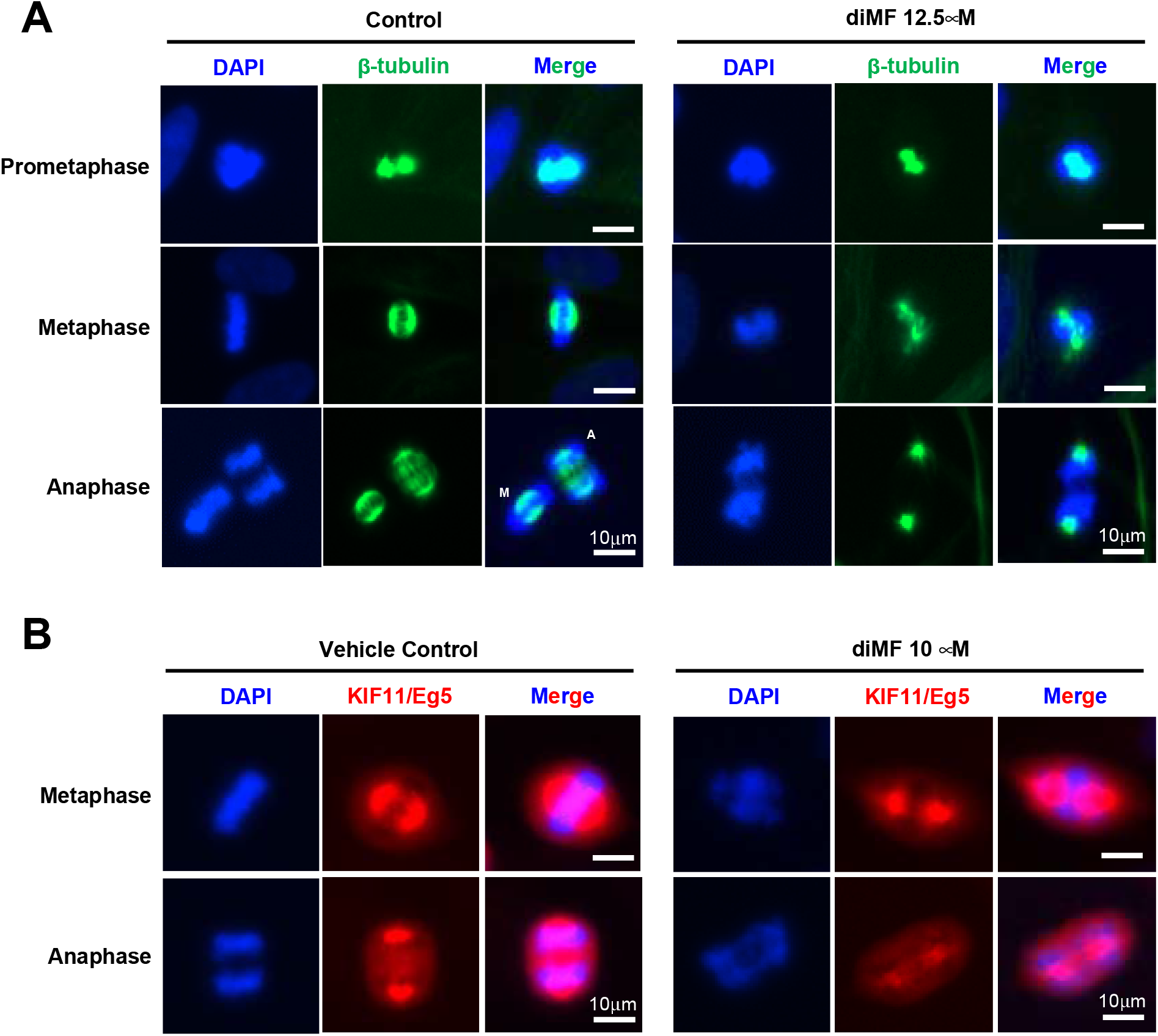
A substandard spindle forms in diMF treated cells. **(A)** RPE-MYC^BCL2^ cells were treated with diMF (12.5 μM) for 6 hours before immunofluorescent staining for β-tubulin to mark microtubules at different stages of mitosis. **(B)** RPE-MYC^BCL2^ cells were treated with diMF (10 μM) for 6 hours before fixation and immunofluorescent staining for the KIF11 (also known as Eg5). DAPI staining in **(A)** and **(B)** was used to visualize DNA. Vehicle control was 0.1% DMSO. All magnification bars equal 10 μm.

### Kinetochore defect and segregation anomalies in diMF treated cells

Next, we examined the integrity of the kinetochores, the protein complexes that connect centromeres to the microtubules of the mitotic spindle. Proper kinetochore assembly supports a robust and geometrically aligned spindle and this, in turn, helps safeguard against chromosome mis-segregation (Magidson et al., 2015). In order to examine kinetochore integrity, we immunohistochemically examined several centromeric markers alongside staining with CREST serum, which harbors autoantibodies from a scleroderma patient that recognize kinetochores (Lopez-Robles et al., 2000; Nishikai et al., 1984). In diMF-arrested, pro-metaphase cells we found diminished fluorescence intensity of the BUB1 kinase. This was consistent across the prometaphase DNA (Figure 3A), even though there was no change in CREST immunofluorescent intensity. BUB1 kinase actively recruits multiple other kinetochore components to enable the efficient segregation of chromosomes (Johnson et al., 2004). This finding suggests diMF induces deficiencies that could impair segregation as early as prometaphase.

**Figure 3:**
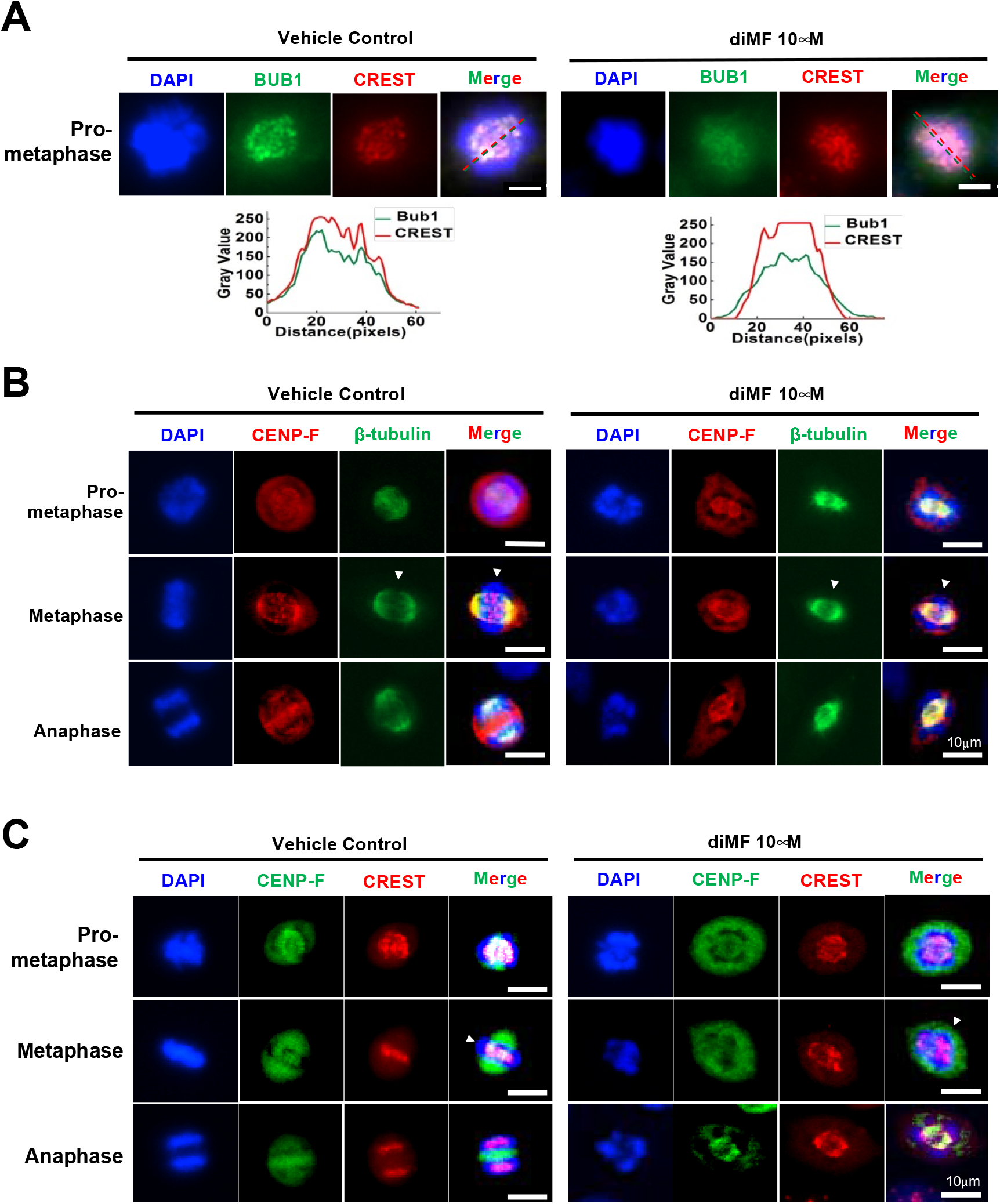
DiMF hinders kinetochore formation. **(A-C)** RPE-MYC^BCL2^ cells were co-stained after 6 hour treatment with diMF (10 μM). **(A)** Cells were stained with an antibody recognizing BUB1 and a CREST serum. A representative cell’s fluorescence intensity was converted to grey value using ImageJ software and the intensity value is shown across the nucleus (dotted lines) for both proteins. **(B)** Cells were co-stained with antibodies recognizing CENP-F and β-tubulin. White arrowheads point to the metaphase plate. **(C)** Cells were co-stained with antibodies recognizing CENP-F and a CREST serum to mark kinetochores (yellow overlap). White arrowheads point to the metaphase plate. Vehicle control for **(A-C)** was 0.1% DMSO. All magnification bars equal 10 μm.

Confirmation of a kinetochore deficiency became apparent with the examination of metaphase cells. The centromeric protein, CENP-F, contains two microtubule-binding domains, and physically associates with dynein motor proteins to stabilize kinetochore-microtubule attachments (Auckland et al., 2020). Co-staining for CENP-F and β-tubulin (Figure 3B) highlighted puncta of CENP-F staining that was present along the metaphase plate in untreated cells, consistent with an association with kinetochores. In diMF treated cells, CENP-F staining was disorganized with no puncta at the metaphase plate. We confirmed that kinetochore staining for CENP-F was absent by co-staining for CENP-F and CREST (Figure 3C). Overlap at the metaphase plate was observed in untreated cells, but not in diMF treated cells. Overall, this suggests defective or delayed formation of the kinetochore complex, which is consistent with the mitotic arrest observed after diMF treatment.

Despite this arrest, a significant number of diMF treated cells did eventually transition through mitosis. We wondered if this was due to failure of the spindle assembly checkpoint. If so, there may be more segregation anomalies present with diMF treatment. We examined cells treated with diMF for 6 hours to detect and catalog mitotic chromosome anomalies using DAPI staining to visualize DNA. Early arrest of cells was dependent on dose of DiMF used as a greater percentage of mitotic cells appeared arrested in prophase/prometaphase with 12.5 μM diMF than with 6.25 μM (Figure 4A). Misalignment of DNA that should have been tightly arranged at the metaphase plate was observed in diMF treated metaphase cells (Figure 4B). Asymmetric segregation of DNA was observed during anaphase (Figure 4C) and in both anaphase and telophase, chromosome bridges persisted (Figure 4D). These observations confirm that errors in chromosome segregation result from diMF treatment of RPE-MYC^BCL2^ cells.

**Figure 4:**
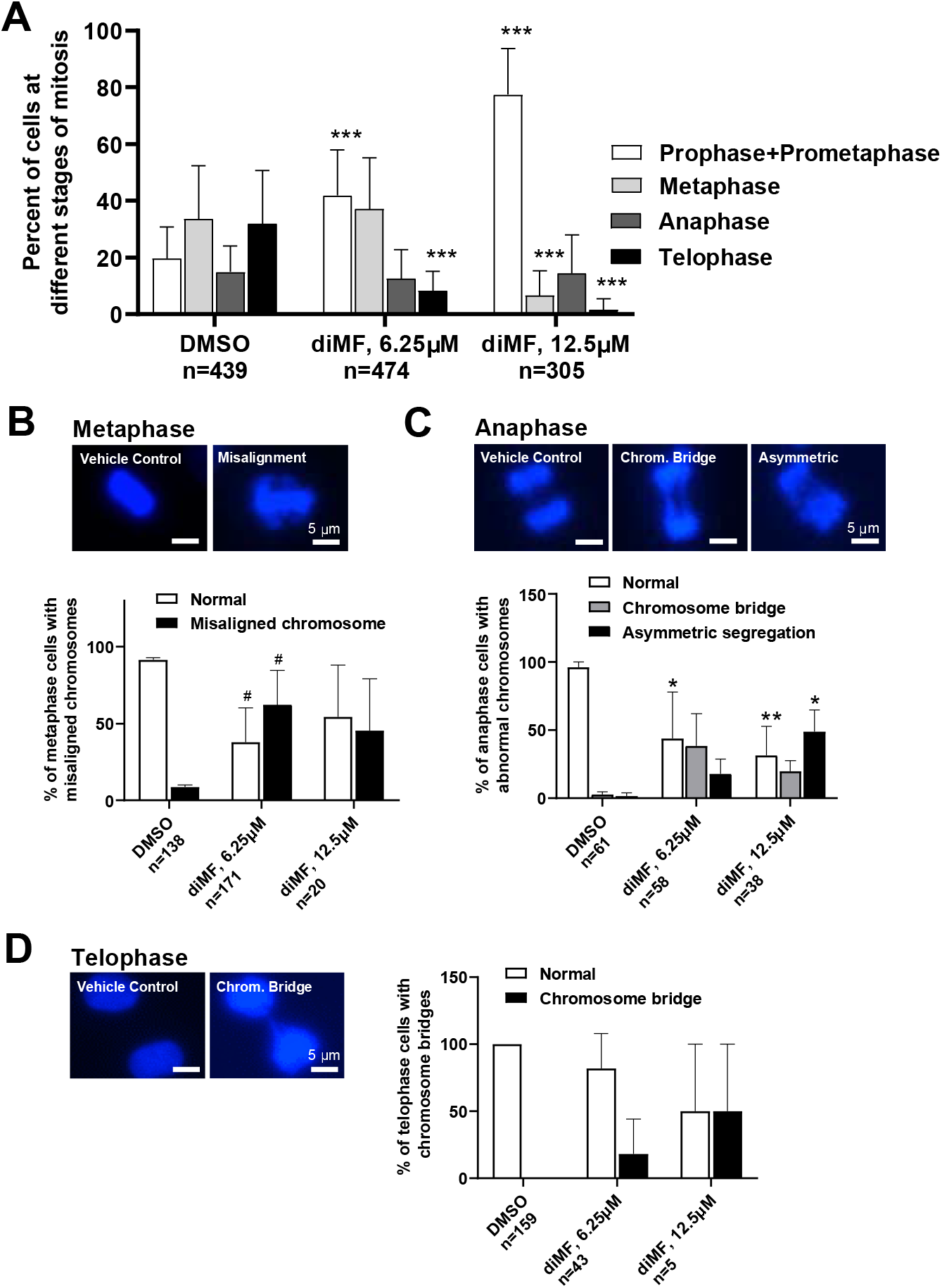
Mitotic chromosome alignment and segregation defects in diMF treated cells. **(A)** Quantification of cells at different stages of mitosis following diMF treatment. RPE-MYC^BCL2^ cells were treated with vehicle control, 6.25 μM or 12.5 μM diMF. DAPI stained nuclei were assessed to determine the stage of mitosis that cells were found at following a 6 hour treatment. Only mitotic cells were quantified and the number of cells counted from 3 replicate plates is shown. **(B)** DAPI stained metaphase cells. Misalignment of chromosomes at the metaphase plate was found and quantified after diMF treatment. **(C)** DAPI stained anaphase cells. Both the presence of chromosome bridges and asymmetric segregations were observed and quantified after diMF treatment. **(D)** DAPI stained telophase cells. Persistent chromosome bridges were observed and quantified in cells after diMF treatment. For **(A-C)**, *** is *p<0.0001*, ** is *p<0.001*, * is *p<0.01* and ^#^ is *p<0.05*. These *p values* are derived from unpaired Student’s t-tests comparing drug to vehicle control (0.1% DMSO) value. All magnification bars equal 5 μm.

### PRC1 is not recruited to the midzone

During anaphase, a set of antiparallel microtubules interdigitate at the equatorial region, or midzone, to form what is known as the central spindle. These antiparallel microtubules interact with proteins that enable antiparallel bundling and movement and serve as an initiation site for the recruitment of factors that participate in cytokinesis (Wadsworth, 2021). Many kinetochore factors are redistributed to the spindle midzone during anaphase (Liu et al., 2020; Wadsworth, 2021), including CENP-F (see anaphase cells in Figure 3B, D for example).

The protein PRC1 binds the antiparallel fibers of the central spindle and its recruitment, along with its motor protein partner, KIF4A, is required for a successful cytokinesis (Kurasawa et al., 2004; Mollinari et al., 2005). PCR1 recruitment is an event that proceeds recruitment of mitotic kinases that are also required to complete cytokinesis (Neef et al., 2007). We examined PRC1 accumulation at the midzone of diMF treated RPE-MYC^BCL2^ cells. PRC1 was absent from the midzone of diMF treated cells. Instead, immunofluorescence for PRC1 was diffuse and without any specific pattern of localization (Figure 5A). This indicates events leading to cytokinetic failure with diMF treatment occur early on in the recruitment of factors regulating the cytokinetic machinery.

**Figure 5:**
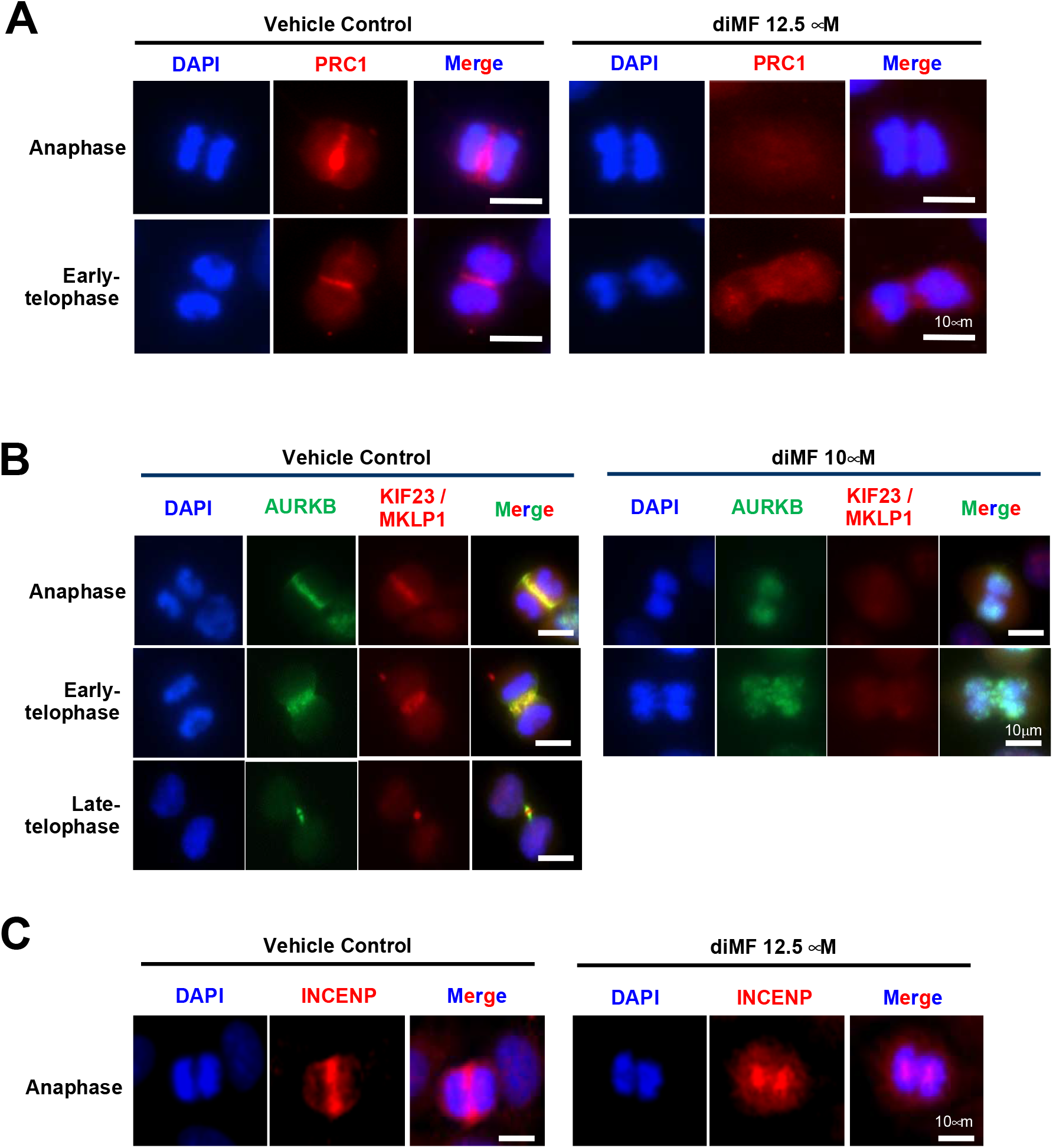
Defective recruitment of central spindle midzone proteins in diMF treated cells. **(A-C)** RPE-MYC^BCL2^ cells were stained after 6 hour treatment with diMF (10-12.5 μM as shown). **(A)** Antibody recognizing the protein regulator of cytokinesis 1 (PRC1) marks the midzone of control cells but not diMF treated cells. **(B)** Co-staining of anaphase and telophase cells with antibodies for AURKB, a component of the chromosomal passenger protein complex (CPP) and KIF23/MKLP1, a component of the centralspindlin complex. Yellow marks overlap at the midzone and midbody of anaphase and telophase cells respectively. **(C)** Antibody staining for a second CPP complex component, INCENP, confirms lack of complex recruitment to the midzone of diMF treated cells. DAPI staining in **(A-C)** was used to visualize DNA. Vehicle control was 0.1% DMSO. All magnification bars equal 10 μm.

### Defective localization of midzone kinases

The complete lack of PRC1 localization at the spindle midzone prompted us to examine the anaphase accumulation of other midzone factors that are important regulators of cytokinesis. Two complexes that bind PRC1 at the midzone are the chromosomal passenger protein (CPP) complex and the centralspindlin complex (Li et al., 2018).

The CPP complex is localized at centromeres during metaphase but it is re-localized and binds PRC1 at the midzone during anaphase and the midbody during telophase to create a gradient of AURKB activity that functions in assembly of the cytokinetic machinery (Minoshima et al., 2003; Ozlu et al., 2010). Knockdown of AURKB or of other components of the CPP results in the formation of polyploid cells (Yang et al., 2010). We observed that the localization of CPP components, AURKB (Figure 5B) and INCENP (Figure 5C), were lacking from the midzone in diMF treated RPE-MYC^BCL2^ cells. The penetrance of this phenotype was examined in human cancer cells and found to be complete. All human cancer cells examined lacked AURKB localization at the midzone following diMF treatment (Figure S4A). This function was specific to diMF as ROCK inhibitors that were not MYC synthetic lethal did not block midzone accumulation of AURKB (Figure S4B).

Targets of AURKB at the midzone include both of the components of the centralspindin complex, the motor protein KIF23 (also MKLP1) (Guse et al., 2005) and the Rho GTPase-activating protein (GAP), called MgcRacGAP (RACGAP1) in vertebrates (Minoshima et al., 2003). Consistently, we found that along with PRC1 and AURKB, the accumulation of KIF23 was also absent from the midzone (Figure 5B), suggesting the centralspindin complex fails to localize here in diMF treated cells.

PRC1 also binds to polo-like kinase 1 (PLK1) to localize this mitotic kinase to the midzone during anaphase. Early in mitosis, PRC1 is a target of cyclin-dependent kinase 1 (CDK1)/cyclin B. By phosphorylating PRC1 at Threonine 470 and 481, CDK1 prevents PRC1 from being a substrate for PLK1 (Neef et al., 2007). As cyclin B activity fades during mitosis, CDK1 no longer targets PRC1 thereby enabling PLK1 to phosphorylate PRC1 and create its own docking site. Immunofluorescent staining for PLK1 in diMF treated cells demonstrated a complete lack of midzone staining in anaphase cells (Figure 6A). PLK1 staining was still found at the centrosomes of diMF treated cells, so this finding is consistent with a PRC1 defect and not a lack of PLK1 catalytic activity.

**Figure 6:**
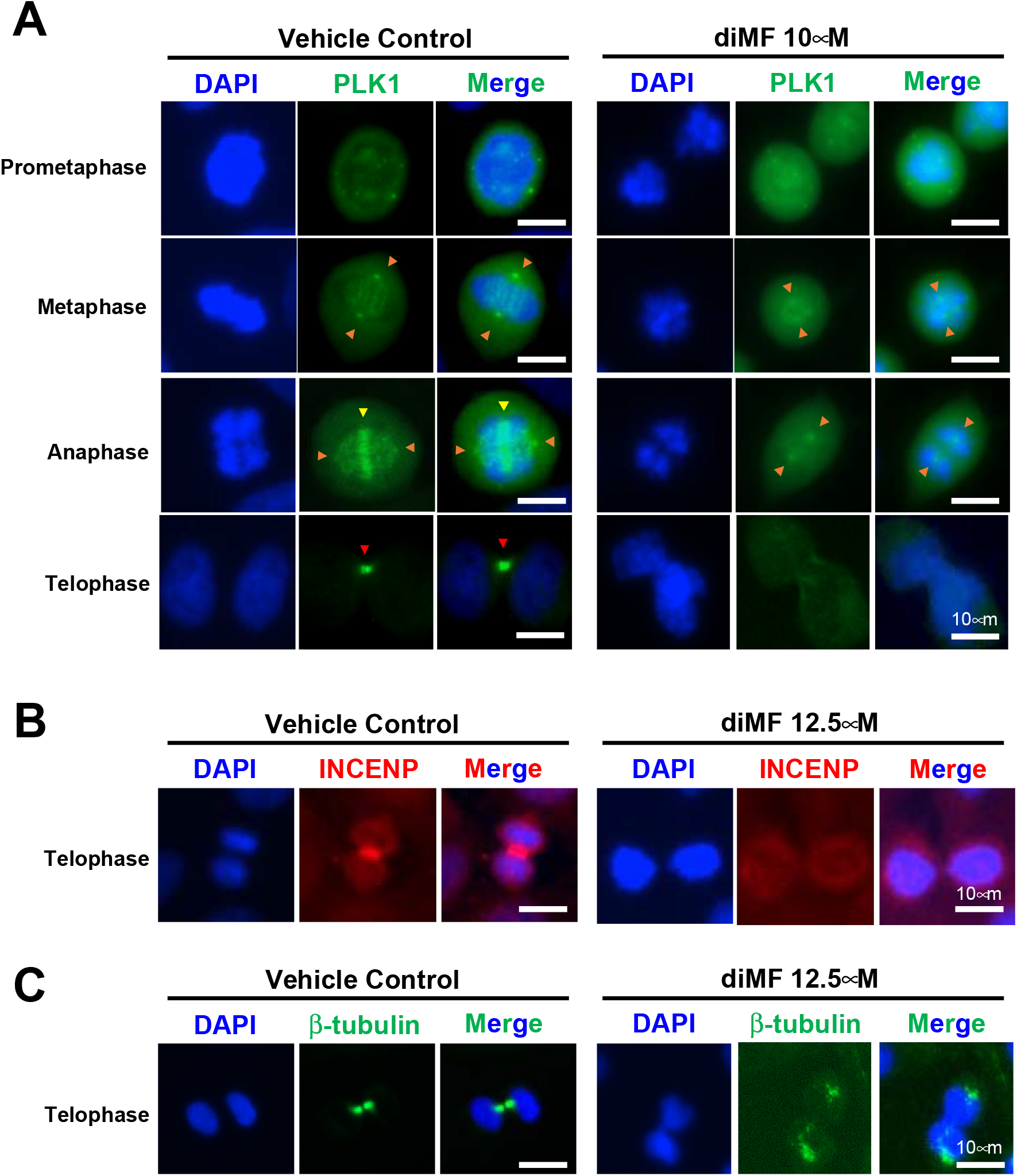
Disruption of midbody formation underlies cytokinetic failure. **(A)** Immunofluorescent staining for polo-like kinase 1 (PLK1) in vehicle control and diMF (101. μM) treated cells. Orange arrowheads mark centrosomes stained with PLK1 antibody. Yellow arrowheads mark midzone staining for PLK1 in anaphase cells. Red arrowheads mark midbody staining for PLK1 in telophase cells. **(B)** Immunofluorescent staining for INCENP in telophase cells treated with diMF (12.5 μM). **(C)** Staining of midbody microtubules with β-tubulin antibody in telophase cells treated with diMF (12.5 μM). In **(A-C)**, DAPI staining was used to mark DNA and assess stage of the cell cycle. Vehicle control was 0.1% DMSO. All magnification bars equal 10 μm.

Midzone defects resulted in complete lack of a midbody in cells that appeared to have progressed to telophase, based on our examination of DAPI stained DNA. Staining for PLK1 (Figure 6A) and the CPP component INCENP (Figure 6B) were not found at the midbody of diMF treated telophase cells, indicating no recovery from deficits observed in anaphase. Examination of β-tubulin (Figure 6C) also produced a complete lack of staining, suggesting the region is not only absent of proteins that direct cytokinesis, but also bereft of microtubules. These observations suggest diMF-induced polyploidy follows from the midzone defects observed and the lack of a midbody in telophase. The components required to direct abscission are spatially lacking.

## Discussion

MYC is a master control factor for programs regulating proliferation, metabolism, angiogenesis and immune escape (Dhanasekaran et al., 2022; Stine et al., 2015). However, as a target for cancer therapy, MYC has long been thought of as an enticing but intractable drug target. As a transcription factor, it is not a suitable target for an inhibitory antibody and it lacks a discernable ATP-binding pocket for high affinity binding of small molecule inhibitors. An alternative approach is to target MYC expressing cells with a drug that is synthetic lethal with MYC. Synthetic lethal drugs attack cellular vulnerabilities introduced as a result of oncogenic changes. Finding and therapeutically targeting these vulnerabilities is very difficult, but synthetic lethal targeting of MYC would enable attack on the many distinct cancer indications driven by MYC while leaving normal tissues, without high MYC expression, relatively untouched.

Previously, we identified dimethylfasudil using a high-throughput screen for compounds that attack RPE cells that ectopically express MYC, but have no effect on cells without MYC (Zhang et al., 2021). Here, we conducted an extensive study of diMF’s dependency on MYC expression for anticancer activity. Our data confirm that anticancer activity correlates with MYC in cultured human cancer cells of varying indications and that the preference persists *in vivo*. In human lung cancer xenografts, activity was highly dependent MYC expression. Our larger study confirms that diMF is synthetic lethal with MYC, although at low micromolar potency in most cellular assays. This may mean diMF is not potent enough to be a clinical candidate itself. However, diMF is a relatively small molecule. It may best serve as a core upon which to build larger analogs that have improved potency and bioavailability. This may be a route to an efficacious MYC synthetic lethal therapy for patients.

DiMF is an analog of clinically used ROCK inhibitors, fasudil (tradename Eril^TM^) and ripasudil (tradename Glanatec^®^). However, cell killing in RPE-MYC and RPE-MYC^BCL2^ cells appears to be through a novel mechanism-of-action, not ROCK inhibition. Analogs used here differ only at the site of one of two methyl groups, one on the hyperpiperazine ring and one on the isoquinoline ring, and they do not recapitulate the killing of MYC expressing cells. Neither does diMF appear to be a spindle toxin. Our immunohistochemical staining for tubulins suggested a reduced spindle tubulin polymer content in diMF-treated cells, but it is unclear if this directly contributes to synthetic lethality. In our experimental RPE cells, paclitaxel, nacodazole and Kif11/Eg5 inhibitors do not recapitulate MYC synthetic lethality (Zhang et al., 2021). Cells without MYC expression are equally susceptible to these drugs with proliferative arrest and cell death, but without widespread polyploid cell induction. The phenotypic response of cells to drugs targeting the spindle is therefore not a match with diMF.

An shRNA screen previously suggested that AURKA might be targeted by diMF (Wen et al., 2012). In addition, AURKB knockdown in RPE-MYC cells (Yang et al., 2010) recapitulates the phenotype induced by diMF (Zhang et al., 2021). This includes the induction of transient cell cycle arrest with mitotic catastrophe-induced apoptosis for a subset of cells and failure to complete cytokinesis for cells that do manage to exit mitosis. Eventually, a lethal autophagy works to eliminates polyploid cells resulting from failed cytokinesis (Zhang et al., 2020a; Zhang et al., 2020b). In addition, either of AURKA (Takahashi et al., 2015) or AURKB (Yang et al., 2010) inhibition favors cell death when the MYC oncoprotein is overexpressed. Despite the phenotypic similarities, diMF does not appear to inhibit the catalytic activity of AURKs and we have now confirmed this in several human cancer cell lines. However, the studies presented here suggest mechanisms that may be just as effective in blocking the mitotic steps that require regulation by mitotic kinases.

DiMF treatment led to the induction of the spindle assembly checkpoint (SAC). The spindle assembly checkpoint (SAC) delays the segregation of chromosomes until proper kinetochore assembly is achieved at all chromosomes. The CPP complex localizes to the kinetochore at early stages of mitosis and, via the kinase activity of AURKB, maintains the SAC through a mechanism that involves tension at the chromosome-spindle attachment site (Carmena et al., 2012; Krenn and Musacchio, 2015). Other kinases also work to maintain kinetochore integrity at the SAC, including the budding uninhibited by benzimidazole 1 (BUB1) protein. BUB1 localizes to the kinetochore very early during mitosis, enabling assembly of kinetochore components, including CENP-F, BUBR1, CENP-E and the essential spindle checkpoint protein, MAD2 (Ciossani et al., 2018; Johnson et al., 2004). We found a deficiency in BUB1 staining as early as the prometaphase arrested cell. In cells that progress further through mitosis, kinetochore integrity was deficient as CENP-F appeared not to co-localize with CREST serum antigen at the metaphase plate. DiMF induced an abundance of asymmetric segregations and chromosome bridges. One might speculate that these resulted from diMF’s effect on kinetochore integrity. However, as mentioned, we also observed reduced tubulin polymer density at the mitotic spindle. We cannot say at this time if kinetochore defects result from underlying spindle deficiencies or are separately induced by diMF, but in theory, both deficiencies could contribute to the mitotic catastrophe and/or segregation defects seen with diMF treatment.

Cytokinetic failure is a prominent phenotype induced by diMF. This can lead to cell loss through the induction of lethal autophagy in polyploid cells, thereby representing a second mode of MYC synthetic lethal cell death (Yang et al., 2010; Zhang et al., 2020a; Zhang et al., 2020b). Cytokinetic failure appears to result from a mis-localization of key factors at the spindle midzone. Notably, we found PRC1 was not recruited to the midzone during anaphase. PRC1 serves as a platform for the localization of key complexes and kinases such as the centralspindlin complex, the CPP complex and PLK1 (Li et al., 2018), so lack of PRC1 may be all that was required to disrupt midzone maturation and abscission. We found loss of the CPP at the midzone in different human cancer cell lines treated with diMF and cells with >4n DNA content were present indicating successive rounds of mitosis without abscission in human cancer cells.

In summary, we observed cell division defects downstream of diMF that were linked with the localization patterns of key regulatory proteins and complexes. Failed recruitment was observed alongside diminished spindle tubulin density prior to and at the time of chromosome segregation and later with absence of β-tubulin from the midbody. The observations presented here can serve as a starting point for the investigation of phenotypes induced by more potent MYC synthetic lethal compounds or for phenotypic screening assays aiming to discover new candidate compounds. We can only speculate at this time that the target of diMF may be one or more of the ATP-dependent motor proteins that are required for proper localization of mitotic kinases. We know of no single motor protein where inhibition might explain all of the phenotypes observed, so we favor the hypothesis that diMF targets multiple motor proteins that regulate spindle dynamics and mitotic kinase localization.

---

**Supplementary Table 1:** Cell lines used to analyze the relationship between MYC expression and diMF sensitivity.

| Cell lines | Primary site/Tissue subtype | Histology | MYC Expression | MYC Analysis Group | Media | Growth properties |
| --- | --- | --- | --- | --- | --- | --- |
| Raji | Haematopoietic/lymphoid tissues | Burkitt's lymphoma | High Myc++ | MYC<sup>High</sup> | RPMI+10%FCS | Suspension |
| Daudi | Haematopoietic/lymphoid tissues | Burkitt's lymphoma | High Myc++ | MYC<sup>High</sup> | RPMI+10%FCS | Suspension |
| HL-60 | Haematopoietic/lymphoid tissues | Acute myeloid leukemia | High Myc++ | MYC<sup>High</sup> | RPMI+10%FCS | Suspension |
| Jurkat | Haematopoietic/lymphoid tissues | Acute T cell leukemia | High Myc++ | MYC<sup>High</sup> | RPMI+10%FCS | Suspension |
| IM-9 | Haematopoietic/lymphoid tissues | Plasma cell myeloma | High Myc++ | MYC<sup>High</sup> | RPMI+10%FCS | Suspension |
| U-937 | Haematopoietic/lymphoid tissues | Histiocytic lymphoma | High Myc++ | MYC<sup>High</sup> | RPMI+10%FCS | Suspension |
| MOLT-4 | Haematopoietic/lymphoid tissues | Acute lymphoblastic leukemia | High Myc+++ | MYC<sup>High</sup> | RPMI+10%FCS | Suspension |
| K-562 | Haematopoietic/lymphoid tissues | Chronic myeloid leukemia | High Myc+++ | MYC<sup>High</sup> | DMEM+10%FCS | Suspension |
| CCRF-CEM | Haematopoietic/lymphoid tissues | Acute lymphoblastic leukemia | High Myc++ | MYC<sup>High</sup> | RPMI+10%FCS | Suspension |
| NCI-BL1672 | Peripheral blood | B lymphoblast; EBV-transformed | Low | MYC<sup>Low</sup> | RPMI+10%FCS | Suspension |
| AG2203 | Autonomic ganglia | Neuroblastoma | Low | MYC<sup>Low</sup> | DMEM+10%FCS | Adherent |
| SH-N-SH | Autonomic ganglia | Neuroblastoma | Low | MYC<sup>Low</sup> | DMEM+10%FCS | Adherent |
| SK-N-AS | Autonomic ganglia | Neuroblastoma | Low | MYC<sup>Low</sup> | DMEM+10%FCS | Adherent |
| SH-SY5Y | Autonomic ganglia | Neuroblastoma | Low | MYC<sup>Low</sup> | MEM+10%FCS | Adherent |
| SK-N-BE2 | Autonomic ganglia | Neuroblastoma | High MycN | MYC<sup>High</sup> | DMEM+10%FCS | Adherent |
| IMR-32 | Autonomic ganglia | Neuroblastoma | High MycN | MYC<sup>High</sup> | MEM+10%FCS | Adherent |
| SK-N-DZ | Autonomic ganglia | Neuroblastoma | High MycN | MYC<sup>High</sup> | DMEM+10%FCS | Adherent |
| NMB | Autonomic ganglia | Neuroblastoma | High MycN | MYC<sup>High</sup> | RPMI+10%FCS | Adherent |
| LAN-1 | Autonomic ganglia | Neuroblastoma | High MycN | MYC<sup>High</sup> | RPMI+10%FCS | Adherent |
| LAN-2 | Autonomic ganglia | Neuroblastoma | High MycN | MYC<sup>High</sup> | RPMI+10%FCS | Adherent |
| LAN-5 | Autonomic ganglia | Neuroblastoma | High MycN | MYC<sup>High</sup> | RPMI+10%FCS | Adherent |
| T98G | Central nervous system/brain | Glioma | Low | MYC<sup>Low</sup> | MEM+10%FCS | Adherent |
| BT-549 | Breast | Ductal carcinoma/papillary | Low | MYC<sup>Low</sup> | RPMI+10%FCS | Adherent |
| BT-474 | Breast | Ductal carcinoma | Low | MYC<sup>Low</sup> | RPMI+10%FCS | Adherent |
| HCC1569 | Breast | Carcinoma | Low | MYC<sup>Low</sup> | RPMI+10%FCS | Adherent |
| HCC1500 | Breast | Breast cancer | High Myc+ | MYC<sup>High</sup> | RPMI+10%FCS | Adherent |
| HCC2157 | Breast | Ductal carcinoma | High Myc++ | MYC<sup>High</sup> | RPMI+10%FCS | Floating aggregates |
| MCF7 | Breast | Carcinoma | High Myc++ | MYC<sup>High</sup> | RPMI+10%FCS | Adherent |
| MDA-MB-157 | Breast | ductal carcinoma/medullary | High Myc++ | MYC<sup>High</sup> | DMEM+10%FCS | Adherent |
| MDA-MB-1751V | Breast | Breast cancer | High Myc+ | MYC<sup>High</sup> | DMEM+10%FCS | Adherent |
| MDA-MB-175VII | Breast | Ductal carcinoma | Low | MYC<sup>Low</sup> | DMEM+10%FCS | Adherent |
| MDA-MB-231 | Breast | Carcinoma | High Myc+ | MYC<sup>High</sup> | DMEM+10%FCS | Adherent |
| MDA-MB-301 | Breast | Breast cancer | Low | MYC<sup>Low</sup> | DMEM+10%FCS | Adherent |
| MDA-MB-415 | Breast | Carcinoma | Low | MYC<sup>Low</sup> | DMEM+10%FCS | Adherent |
| MDA-MB-436 | Breast | Breast cancer | High Myc++ | MYC<sup>High</sup> | L-15+10%FCS | Adherent |
| MDA-MB-468 | Breast | Carcinoma | High Myc++ | MYC<sup>High</sup> | L-15+10%FCS | Adherent |
| T47D | Breast | Ductal carcinoma | Low | MYC<sup>Low</sup> | RPMI+10%FCS | Adherent |
| ZR-751 | Breast | Breast cancer | Low | MYC<sup>Low</sup> | RPMI+10%FCS | Adherent |
| NCI-H146 | Lung | Small cell carcinoma | Low | MYC<sup>Low</sup> | RPMI+10%FCS | Floating aggregates |
| NCI-H128 | Lung | Small cell carcinoma | Low | MYC<sup>Low</sup> | RPMI+10%FCS | Floating aggregates |
| NCI-H1299 | Lung | Large cell carcinoma | High Myc++ | MYC<sup>High</sup> | RPMI+10%FCS | Adherent |
| NCI-H209 | Lung | Small cell carcinoma | High L-Myc | MYC<sup>High</sup> | RPMI+10%FCS | Floating aggregates |
| NCI-H2170 | Lung | squamous cell carcinoma | High Myc+++ | MYC<sup>High</sup> | RPMI+10%FCS | Adherent |
| NCI-H2171 | Lung | Small cell carcinoma | High Myc+++ | MYC<sup>High</sup> | RPMI+10%FCS | Floating aggregates |
| NCI-H2199 | Lung | Lung cancer (NSCLC) | Low | MYC<sup>Low</sup> | RPMI+10%FCS | Adherent |
| NCI-H2195 | Lung | Small cell carcinoma | Low | MYC<sup>Low</sup> | DMEM+10%FCS | Adherent |
| NCI-H211 | Lung | Small cell carcinoma | Low | MYC<sup>Low</sup> | RPMI+10%FCS | Adherent |
| NCI-H23 | Lung | non_small_cell_carcinoma | High Myc++ | MYC<sup>High</sup> | RPMI+10%FCS | Adherent |
| NCI-N417 | Lung | Small cell carcinoma | High Myc+++ | MYC<sup>High</sup> | DMEM+10%FCS | Floating aggregates |
| NCI-H510A | Lung | Small cell carcinoma | High L-Myc | MYC<sup>High</sup> | RPMI+10%FCS | Floating aggregates |
| NCI-H520 | Lung | squamous cell carcinoma | Low | MYC<sup>Low</sup> | RPMI+10%FCS | Adherent |
| NCI-H524 | Lung | Small cell carcinoma | High Myc+++ | MYC<sup>High</sup> | RPMI+10%FCS | Floating aggregates |
| NCI-H526 | Lung | Small cell carcinoma | High MycN | MYC<sup>High</sup> | RPMI+10%FCS | Floating aggregates |
| NCI-H82 | Lung | Small cell carcinoma | High Myc+++ | MYC<sup>High</sup> | RPMI+10%FCS | Suspension |
| NCI-H841 | Lung | Small cell carcinoma | High Myc++ | MYC<sup>High</sup> | RPMI+10%FCS | Adherent |
| A549 | Lung | Carcinoma | Low | MYC<sup>Low</sup> | RPMI+10%FCS | Adherent |
| Calu-6 | Lung | Adenocarcinoma | High Myc++ | MYC<sup>High</sup> | RPMI+10%FCS | Adherent |
| DU-145 | Prostate | Adenocarcinoma | High Myc++ | MYC<sup>High</sup> | DMEM+10%FCS | Adherent |
| LNCaP | Prostate | Adenocarcinoma | High Myc++ | MYC<sup>High</sup> | RPMI+10%FCS | Adherent |
| VCaP | Prostate | Adenocarcinoma | High Myc++ | MYC<sup>High</sup> | DMEM+10%FCS | Adherent |
| HT-29 | Large intestine/Colon | Carcinoma | High Myc++ | MYC<sup>High</sup> | McCoy's 5A +10%FCS | Adherent |
| HCT-116 | Large intestine/Colon | Carcinoma | High Myc++ | MYC<sup>High</sup> | McCoy's 5A +10%FCS | Adherent |
| SW480 | Large intestine/Colon | Adenocarcinoma | High Myc++ | MYC<sup>High</sup> | L15+10%FCS | Adherent |
| SaOs-2 | Bone | Osteosarcoma | High Myc++ | MYC<sup>High</sup> | DMEM+10%FCS | Adherent |
| U2OS | Bone/Tibia | Osteosarcoma | High Myc++ | MYC<sup>High</sup> | DMEM+10%FCS | Adherent |
| Hela | Cervix | Carcinoma | High Myc++ | MYC<sup>High</sup> | DMEM+10%FCS | Adherent |
| C-33-A | Cervix | Carcinoma | High Myc++ | MYC<sup>High</sup> | MEM+10%FCS | Adherent |
| ACHN | Kidney | Renal cell carcinoma | High Myc++ | MYC<sup>High</sup> | MEM+10%FCS | Adherent |
| TUWI | Kidney | Wilms' tumor | High Myc++ | MYC<sup>High</sup> | DMEM+10%FCS | Adherent |
| SK-N-EP1 | Kidney | Wilms' tumor | High Myc+++ | MYC<sup>High</sup> | McCoy's 5A +10%FCS | Adherent |
| HT-1080 | Soft tissue | Fibrosarcoma | Low | MYC<sup>Low</sup> | MEM+10%FCS | Adherent |
| MDA-MB-435 | Breast | Carcinoma | High Myc++ | MYC<sup>High</sup> | DMEM+10%FCS | Adherent |
| A431 | Skin | squamous cell carcinoma | High Myc+ | MYC<sup>High</sup> | DMEM+10%FCS | Adherent |
| T-24 | Urinary tract/bladder | Transmonal cell carcinoma | Low | MYC<sup>Low</sup> | McCoy's 5A +10%FCS | Adherent |

**Supplementary Figure 1:**
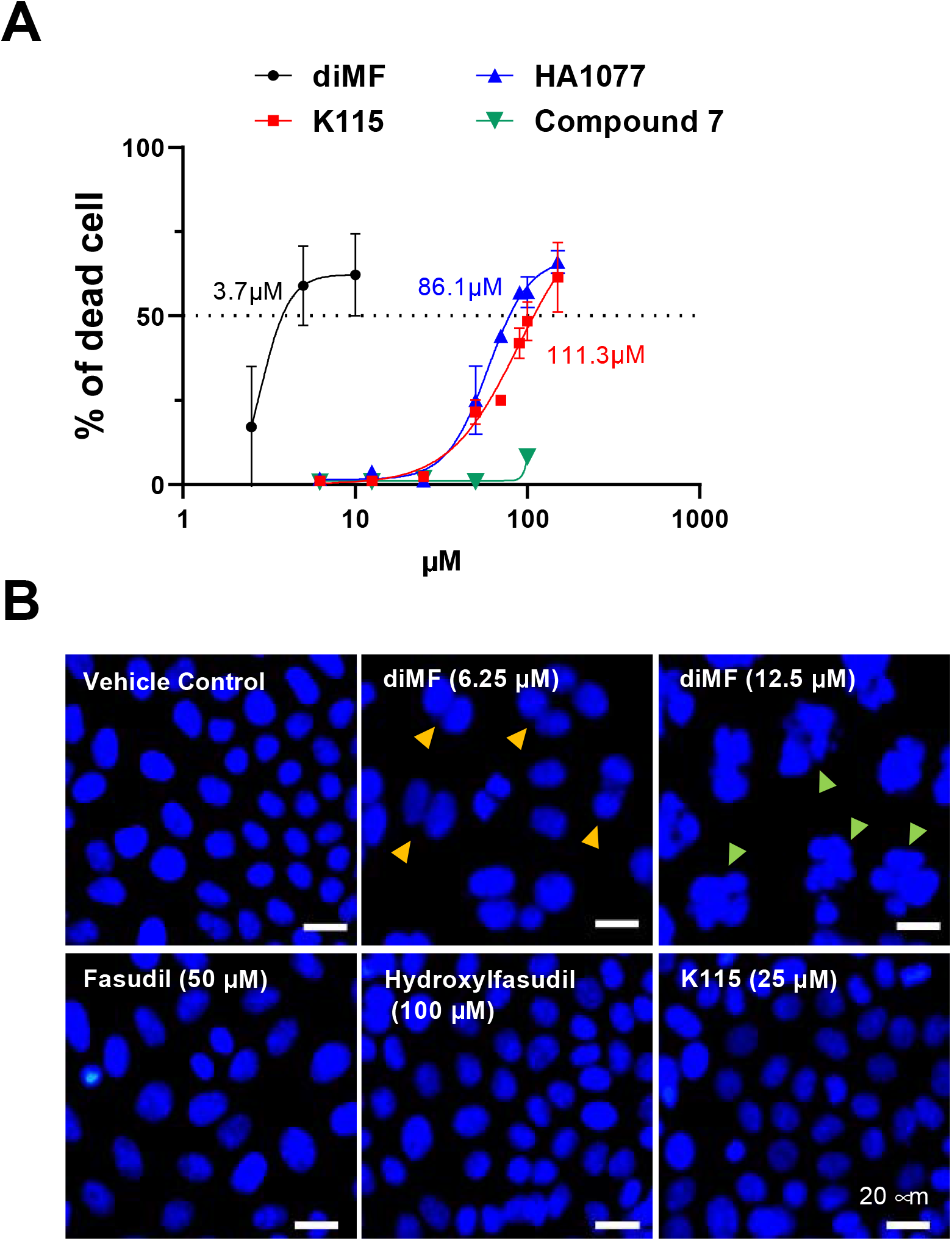
ROCK inhibitors that are structurally related to diMF do not induce mitotic catastrophe or polyploidy in RPE-MYC^BCL2^ cells. **(A)** Concentration-response curves for diMF, K115, HA1077 and compound 7, after a 48 hour exposure. EC_50_ values were estimated with curve fitting using Prism software. For each drug concentration, the percent of dead cells to total cells was averaged from three separate experiments with 3 replicates in each (n=9), using the trypan blue exclusion assay. **(B)** DAPI staining of nuclei of RPE-MYC^BCL2^ cells after 48 hours of drug exposure. Yellow arrowheads point to bi-nucleate cells. Green arrowheads point to multinucleate polyploid cells. All magnification bars equal 20 μm.

**Supplementary Figure 2:**
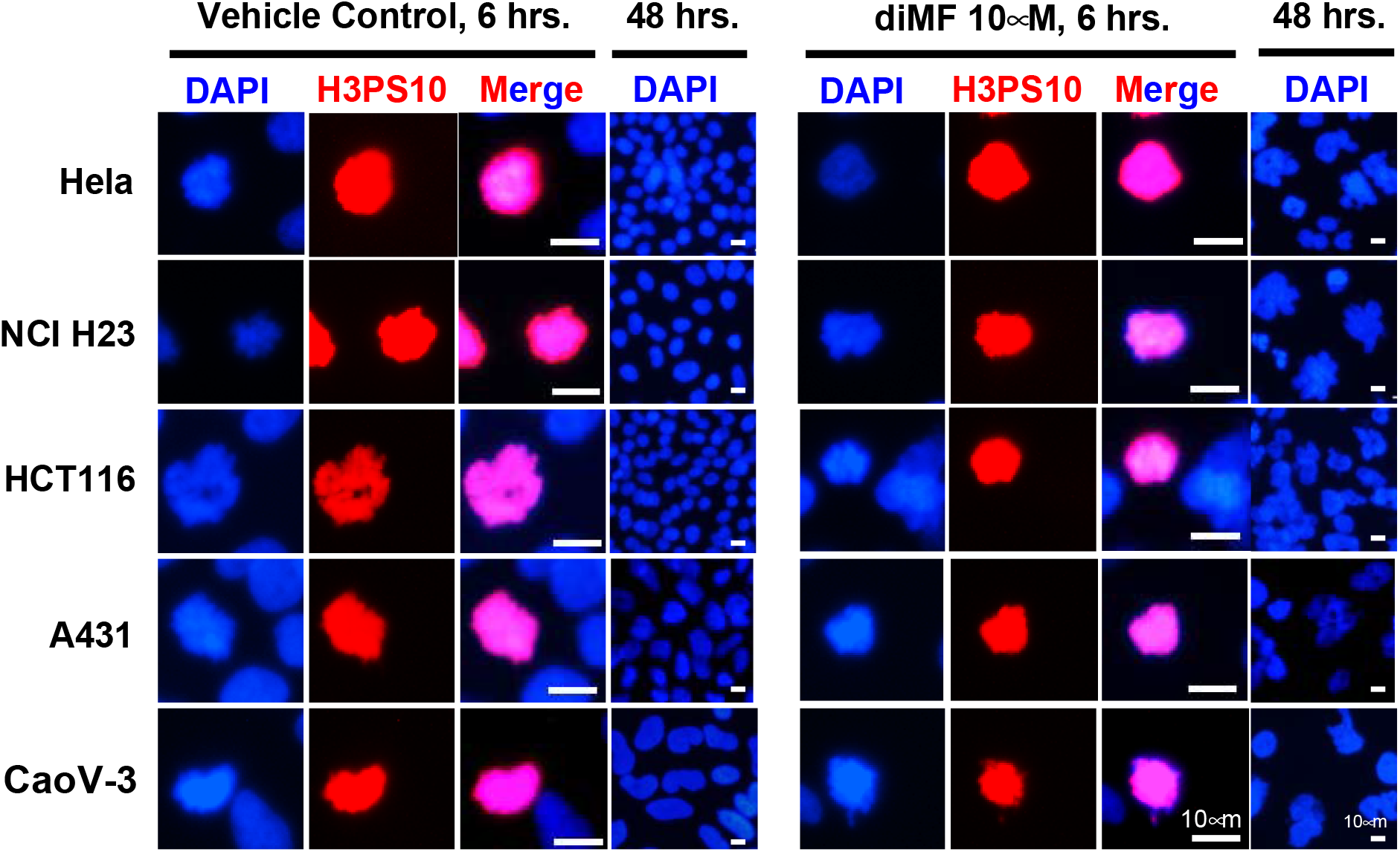
AURKB activity is not diminished by diMF in cells arrested in prometaphase. The human cancer cell lines indicated were treated with diMF (10 μM) for 6 hours before fixation and immunofluorescent staining for histone 3 phosphorylated at serine 10. Individual prometaphase stage cells are shown. Induction of polyploidy in these cell lines was confirmed with DAPI staining after 48 hours to visualize multinucleate cells. Vehicle control was 0.1% DMSO. All magnification bars equal 10 μm.

**Supplementary Figure 3:**
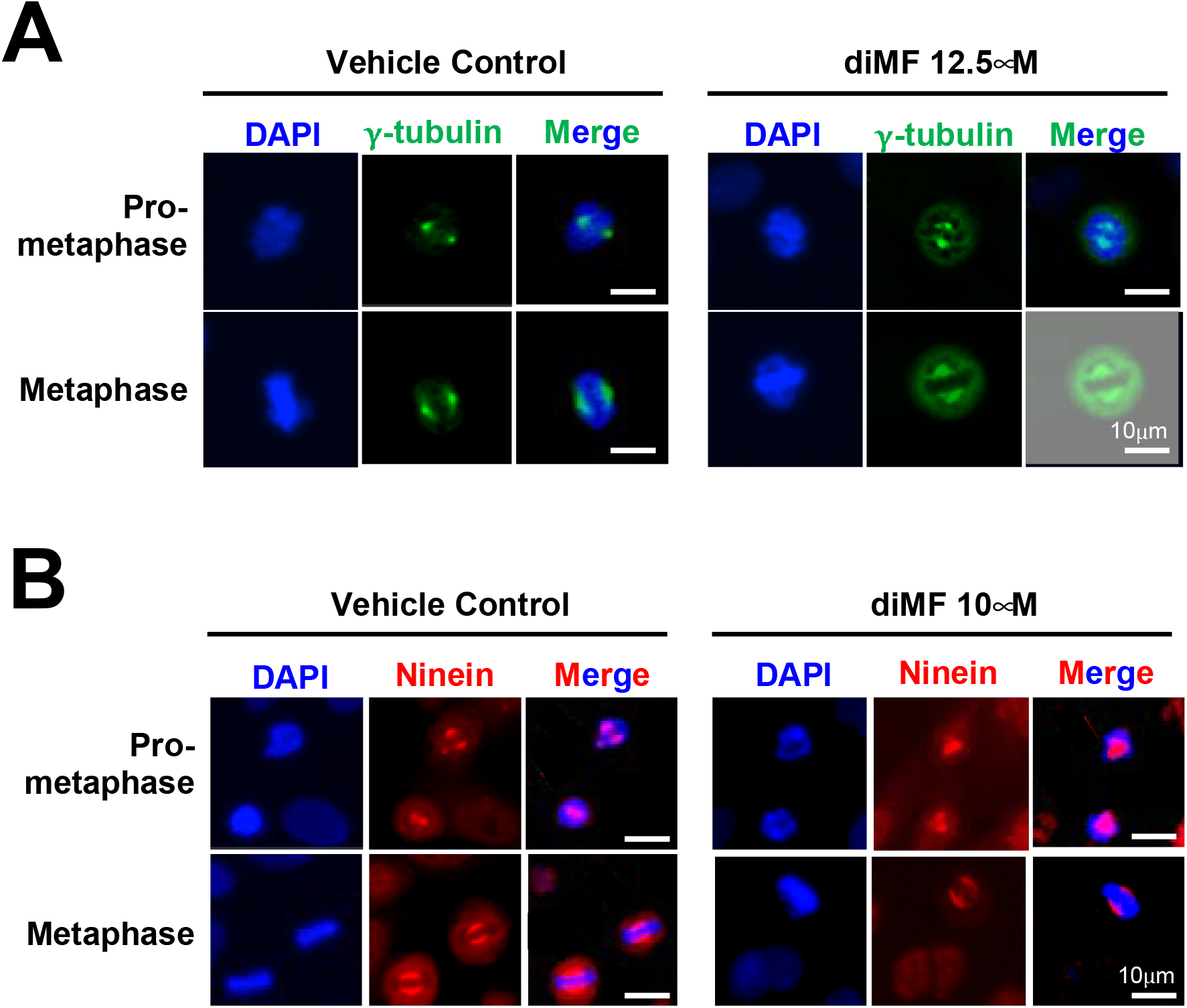
DiMF does not induce centrosome amplification in RPE-MYC^BCL2^ cells. **(A)** RPE-MYC^BCL2^ cells were treated with diMF (12.5 μM) for 6 hours before fixation and immunofluorescent staining for γ-tubulin to mark centrosome location in the dividing cell. **(B)** RPE-MYC^BCL2^ cells were treated with diMF (10 μM) for 6 hours before fixation and immunofluorescent staining for Ninein, a component of the pericentriolar material. DAPI staining in **(A)** and **(B)** was used to visualize DNA. Vehicle control was 0.1% DMSO. All magnification bars equal 10 μm.

**Supplementary Figure 4:**
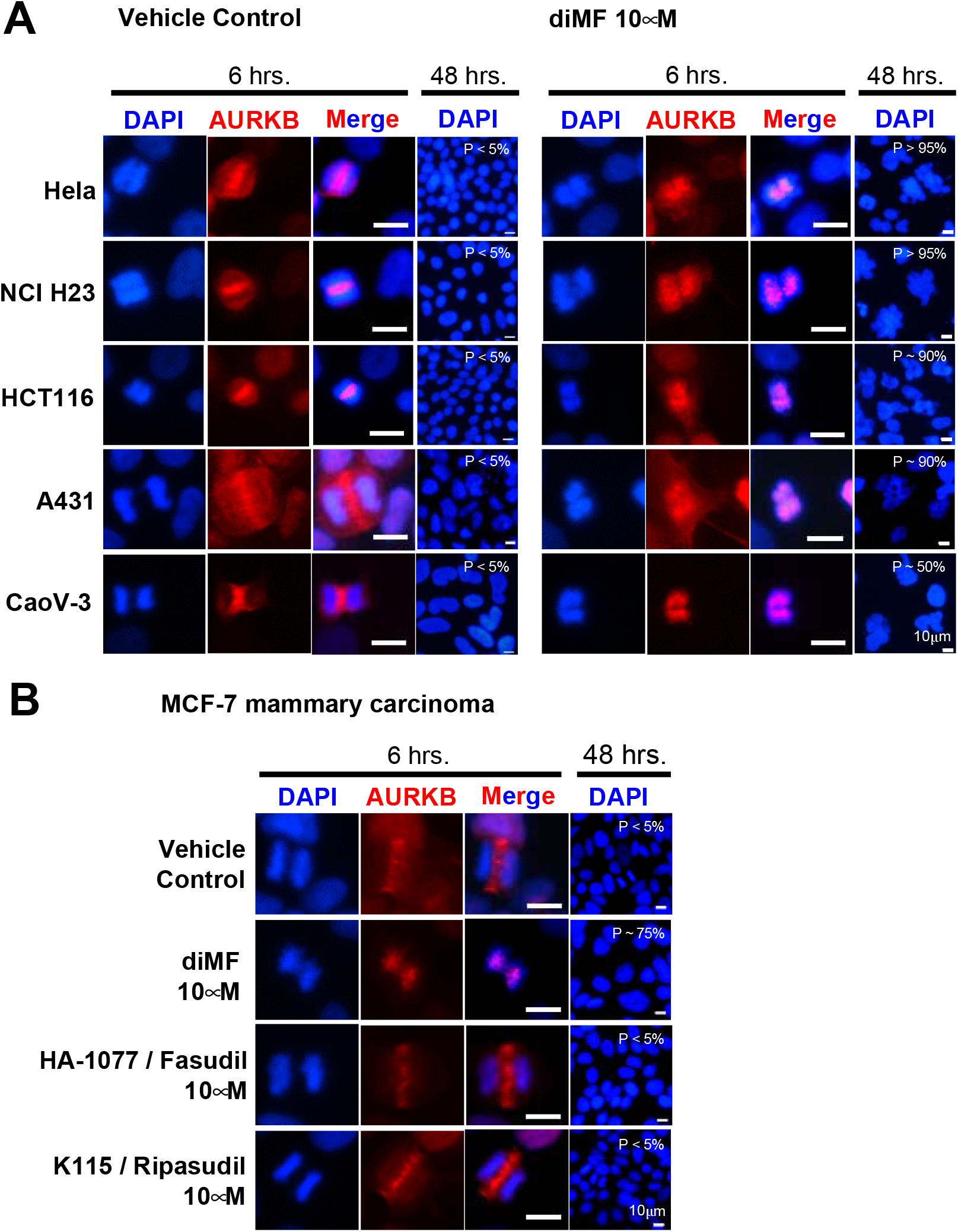
Exclusion of the chromosomal passenger protein complex from the spindle midzone. **(A)** Immunofluorescent staining for AURKB marks the midzone of anaphase cells treated with vehicle control, but it is excluded from the midzone following diMF (10 μM) treatment. **(B)** The mammary cell line MCF7 was treated with the indicated compounds. Cells were stained for AURKB to assess recruitment to the midzone of anaphase cells. In **(A, B)**, DAPI staining was used to mark DNA and assess the stage of the cell cycle and number of nuclei per cell. Induction of polyploidy was confirmed in each human cancer cell line assayed after a 48 hour treatment (P = polyploidy). Vehicle control was 0.1% DMSO. All magnification bars equal 10 μm.

